# IFN-α and 5-Aza-2’-deoxycytidine enhance the anti-tumor efficacy of a dendritic-cell targeting MIP3a-Gp100-Trp2 DNA vaccine by affecting T-cell recruitment and tumor microenvironment gene expression

**DOI:** 10.1101/531616

**Authors:** James T. Gordy, Kun Luo, Richard B. Markham

**Affiliations:** The W. Harry Feinstone Department of Molecular Microbiology and Immunology, Johns Hopkins Bloomberg School of Public Health, Baltimore, MD 21205

## Abstract

**Background:** The chemokine MIP-3α (CCL20) binds to CCR6 found on immature dendritic cells. DNA vaccines fusing MIP-3α to melanoma-associated antigens have shown improved efficacy and immunogenicity in the B16F10 model. To optimize the therapy, our laboratory has added agents designed to overcome immunoregulatory mechanisms of the tumor microenvironment. Here, we report that the combination of type-I interferon therapy (IFNα) with 5-Aza-2’-deoxycitidine (Aza) profoundly enhanced the therapeutic anti-melanoma efficacy of a MIP-3α-Gp100-Trp2 DNA vaccine.

**Methods:** The current studies utilize the B16F10 syngeneic mouse melanoma model. The vaccine is administered intramuscularly (i.m.) followed by i.m. electroporation. Vaccinations are given thrice at one-week intervals beginning day 5 with CpG adjuvant given two days later as noted. Aza is given i.p. at 1mg/kg on days 5 and 12. IFNα therapy is given in a series of one high followed by three low doses, beginning on days 5 and 12. Tumor sizes, growth, and survival were all assessed. Tumor microenvironment gene expression levels were explored by qRT-PCR. Tumor-infiltrating lymphocytes (TILs) were assessed by stimulating the purified lymphocyte fraction of tumors with vaccine antigens followed by intracellular cytokine staining flow cytometry.

**Results:** We demonstrate that the addition of IFNα and Aza treatments to mice vaccinated with the MIP-3α-Gp100-Trp2 vaccine has led to significantly reduced tumor burden and overall increases in mouse survival, increasing median survival by 39% over vaccine and 86% over controls. Importantly, this increase in efficacy was dependent on the presence of all three components: vaccine, IFNα, and Aza. The addition of Aza and IFNα to the vaccine increased T-cell tumor infiltration and altered the proportion of CD8+T-cells. Also, IFNα and vaccine induced durable changes in IFNα-stimulated gene transcription.

**Conclusions:** Efficient targeting of antigen to immature dendritic cells with a chemokine-fusion vaccine offers a potential alternative approach to classic and dendritic cell based vaccines currently undergoing clinical investigation. Combining this approach with IFNα and Aza combination treatment significantly improved vaccine efficacy, with efficacy correlating with changes in TILs and in IFNα -stimulated gene expression. Further potential therapy optimization currently undergoing investigation offers promise for this line of investigation to become a novel melanoma therapy.

## BACKGROUND

The remarkable efficacy of immunologic interventions in controlling certain cancers has prompted renewed interest in the potential of cancer vaccines as one arm of the immunologic approach. A focus of this interest has been identifying more effective adjuvants, which, among other properties, would attract more immune cells to the site of vaccine exposure. However, only 5-10% of inflammatory cells attracted to vaccination sites by adjuvants actually play a role in the initiation of the immune response^1^. To more efficiently direct vaccine antigen to the dendritic cells critical to the initiation of the adaptive immune response, we have been studying the effect of genetically fusing a vaccine antigen to the chemokine macrophage inflammatory protein 3 alpha (MIP3α), also known as chemokine (C-C motif) ligand 20 (CCL20), which targets the vaccine antigen to the C-C chemokine receptor type 6 (CCR6) on immature dendritic cells (iDC) ^2,3^.

The simple addition of this small molecule to the vaccine construct has been shown to enhance immunogenicity and efficacy over antigen-only vaccinations in melanoma^2,4,5^, lymphoma^6^, and malaria model systems^7,8^. A therapeutic melanoma DNA vaccine in which MIP3α is fused to amino acids 25-235 of the human melanoma-associated antigen gp100 proved superior compared to vaccines comprised of defective MIP3α (D-MIP3α) fused to gp100 or MIP3α fused to an irrelevant antigen^4^.

Therapeutic cancer vaccines have historically failed as a monotherapy against established and progressing tumors, but recent studies have highlighted the potential of novel vaccine platforms to be effective when used in combination with other cancer therapies, especially other immunotherapies (reviewed in ^9^). To address immunosuppressive factors in the tumor microenvironment, we examined the effect in the mouse melanoma model of neutralizing IL-10 used in combination with the MIP3α-gp100 vaccine^5^. This approach generated significant, but moderate improvement in survival. Importantly the observed improvement in survival was shown to be mediated by enhanced type 1 interferon activity^5^.

In the current study, we have explored the potential importance of Type I interferons in enhancing the effect of the vaccine-generated immune response. For this purpose, we have added to the therapeutic regimen direct administration of recombinant IFN-α, the type I IFN-stimulating adjuvant CpG, and an inhibitor of methyl transferase activity, as DNA methylation has been shown to attenuate the activity of interferon-stimulated genes (ISG’s) ^10–12^. Also because of evidence from studies supporting the use of multiple antigens in cancer vaccines^13–15^, we have added the melanoma antigen tyrosinase-related protein 2 (Trp2) to our original gp100 vaccine construct. The results of our study indicate that this combined regimen effectively doubles survival time compared to the negative control in this melanoma model system.

## METHODS

### Animals and Tumor Model

5-6 week old female C57BL/6 (H-2b) mice were purchased from Charles River Laboratories (Wilmington, MA) and maintained in a pathogen-free micro-isolation facility in accordance with the National Institutes of Health guidelines for the humane use of laboratory animals. All experimental procedures involving mice were approved by the IACUC of the Johns Hopkins University (Protocol MO16H147). B16F10 mouse melanoma cells were a generous gift from Dr. Jonathan Schneck (Johns Hopkins School of Medicine, Baltimore, MD). B16F10 melanoma cells would be cultured from frozen stock for at least three days and no more than ten passages under sterile conditions utilizing complete growth media (Dulbecco’s Modified Eagles Medium [DMEM+L-glutamine, L glucose, and sodium pyruvate; Corning™ cellgro™, Corning, NY]; 10% Fetal Bovine Serum [FBS, Corning™, Corning, NY]; 0.1% gentamycin [Quality Biological, Gaithersburg, MD]; 2% penicillin/streptomycin [Corning™ cellgro™, Corning, NY]; and 1% non-essential amino acids [Gibco™, Life Technologies, Carlsbad, CA]). Cells were passaged utilizing 0.25% Trypsin (Quality Biological, Gaithersburg, MD). Prior to challenge, cells would be assessed by Gibco™ Trypan Blue solution 0.4% (Life Technologies, Carlsbad, CA), ensuring cell viability ≥95%. 6-8 week old mice were challenged in the left flank subcutaneously with a lethal dose of B16F10 melanoma (5×10^4^ cells in sterile 1xHanks Balanced Salt Solution [Gibco™, Life Technologies, Carlsbad, CA]). Tumor size was recorded as square mm, representing tumor length × width (opposing axes) measured by calipers every 1-3 days. Mice were kept in survival studies until one of the following occurred: mouse death, tumor diameter eclipsing 20mm, significant lethargy, or extensive tumor necrosis resulting in excessive bleeding.

### Plasmid Design

The original plasmid encoded the MIP3α-hgp100 fusion sequence, where the antigen includes amino acids 25-235 of human gp100, as described^4^. A vaccine of mouse MIP3α fused to the combined antigens construct consisting of the hgp100 construct and mouse tyrosinase-related protein 2 (Trp2, amino acids 170-269) was created. The region of Trp2 included a 5’ spacer region (amino acids MEFNDAQAPKSLEA) and was flanked by XbaI restriction sites. This construct was synthesized by Genscript Biotech Corp (Piscataway Township, NJ) in the pUC57 cloning vector. Using standard cloning techniques, the spacer-Trp2 sequence was cloned from pUC57 to downstream of MIP3α-gp100 and also to dMIP3α-gp100^1^using the XbaI restriction enzyme (New England Biolabs, Ipswich, MA) to create MIP3α-Gp100-Trp2 and dMIP3α-Gp100-Trp2 vaccines. MIP3α-Trp2 vaccine was created by using the following primers [F: 5’-ctcgagagtctcgaagctgggctggt-3’ and R: 5’-ctgttcttctgcggatctctctagagtcg −3’] to PCR amplify the Trp2 region from the MIP3α-Gp100-Trp2 plasmid, incorporating an xhoI restriction site at the 5’ end of the construct downstream of extant xbaI site. PCR amplification was performed using Taq DNA Polymerase with Standard Taq Buffer according to the manufacturer’s protocol (New England Biolabs, Ipswich, MA). The PCR product was inserted into pCR™ 2.1-TOPO® TA Cloning® plasmid according to the manufacturer’s protocol (Invitrogen™ ThermoFisher Scientific, Waltham, MA). Utilizing standard cloning techniques, xbaI and xhoI enzymes (New England Biolabs, Ipswich, MA) were used to clone the Trp2 sequence from the pCR™ 2.1-TOPO® TA Cloning® plasmid into the MIP3α-gp100 vaccine plasmid, replacing the gp100 with Trp2 to create a MIP3α-Trp2 vaccine.

### Vaccinations and Therapeutics

Vaccination plasmids were extracted from *E. coli* using Qiagen^®^ (Germantown, MD) EndoFree^®^ Plasmid Maxi, Mega, and Giga Kits and were diluted with endotoxin-free 1xPBS. Vaccine DNA purity, quality, and quantity were verified by gel electrophoresis, restriction enzyme analysis, Nanodrop^®^ spectrophotometry, and insert sequencing (JHMI Synthesis and Sequencing Facility, Baltimore, MD). Mock vaccinations were comprised of endotoxin-free PBS only. DNA injections were administered into the hind leg tibialis muscle. Immediately following injection, the muscle was pulsed using an ECM 830 Electro Square Porator™ with 2-Needle Array™ Electrode (BTX Harvard Apparatus^®^; Holliston, MA) under the following parameters: 106V; 20ms pulse length; 200ms pulse interval; 8 total pulses. Vaccinations of 50ug/dose were delivered at days noted in figure legends. If included in the regimen, 50µg ODN2395 Type C CpG (Innaxon LPS Biosciences, Tewkesbury, UK) was administered two days after vaccination intramuscularly into vaccinated muscle. Recombinant Mouse Interferon Alpha-A (IFNα, R&D Systems, Inc. Minneapolis, MN) was administered either intratumorally or intramuscularly, as indicated. If intramuscular, it was administered into the tibialis muscle of the leg that was not administered vaccine. High doses of IFNα were 10,000 units per dose and low doses were 1,000 units per dose. InSolution™ 5 Aza 2’-deoxycytidine (Aza, CalBiochem®, MilliporeSigma, Burlington, MA) was administered interperitoneally at 1mg/kg in 50µl, at approximately 20µg/mouse.

### Extraction of Splenocytes and TILs

Spleen and tumor cell suspensions were prepared by grinding sterile excised tissue between the frosted ends of microscope slides and then passing the tissue through a sterile 70 µM mesh (Westnet, Inc. Canton, MA). Splenocytes were processed by lysing red blood cells according to manufacturer’s protocol (ACK lysing buffer, Quality Biological, Gaithersburg, MD) and washing with sterile PBS. For tumor lymphocyte analysis, tumor lysate was washed with sterile PBS, and the mononuclear cell fraction (including TILs) was enriched by Lympholyte^®^-M Cell Separation Media (Cedarlane^®^, Burlington, NC) according to the manufacturer’s protocol. Tissues/cells were kept on ice or at 4°C at all points possible. Single cell suspensions were either stimulated immediately or left at 4°C overnight.

### Intracellular Cytokine Staining and Flow Cytometry

Splenocytes or enriched TILs were seeded onto Falcon^®^ Multiwell 24-well tissue culture treated plates (Corning, Inc.; Corning, NY) at approximately 1×10^6^ cells per well (or all cells if total is less) and stimulated for 3-5 hours at 37°C with equal concentrations of known immunodominant peptides gp100_25-33_ (KVPRNQDWL; JHU School of Medicine Synthesis & Sequencing Facility; Baltimore, MD) and Trp2_180-188_ (SVYDFFVWL; Anaspec Inc. Fremont, CA) or with control HA peptide (YPYDVPDYA; JHU School of Medicine Synthesis & Sequencing Facility; Baltimore, MD) for a total of 20µg of peptide per sample. Peptide(s) were combined with Protein Transport Inhibitor Cocktail and costimulatory anti-CD28 and anti-CD49d agonizing antibodies (eBioscience, Inc. San Diego, Ca). Assay positive controls were stimulated with Cell Stimulation Cocktail and Protein Transport Inhibitor Cocktail (eBioscience, Inc. San Diego, Ca). Cells were collected, washed, fixed, permeabilized, and stained using standard laboratory protocols for intracellular staining. Fixation and permeabilization buffers from BD Cytofix/Cytoperm™ Kit (BD Biosciences, San Jose, CA) were used. Stains utilized were the following anti-mouse mAbs: PercPCy5.5 conjugated anti-CD3, APC-IFNγ, FITC-CD8, PE-CD4, PECy7-TNFα, (eBioscience, Inc. San Diego, CA), FITC-CD8, and Live/Dead Near-IR (Invitrogen by Thermo Fisher Scientific, Carlsbad, CA). The Attune™ NxT (Thermo Fisher Scientific, Waltham, MA) flow cytometer was utilized. Flow data were analyzed by FlowJo Software (FlowJo, LLC Ashland, OR). Total cell count estimation was back-calculated from volume utilized by cytometer to create a cell concentration value that could be applied to the total volume of the sample.

### RNA Extraction and qRT-PCR

Mice were sacrificed and portions of tumor weighing less than 100mg were harvested. Tumor was minced as finely as possible, added to 1ml Trizol® (Ambion® by Life Technologies, Carlsbad, Ca), and then homogenized by the Fisher Scientific™ PowerGen 125 (Thermo Fisher Scientific, Waltham, MA). RNA was extracted utilizing the manufacturer’s protocol and including a 75% ethanol wash step. The pellet was air dried and resuspended in nuclease-free water. The cDNA Reverse Transcription reaction was performed with 1µg extracted RNA using the High Capacity cDNA Reverse Transcription Kit with random primers (Applied Biosystems™ by Thermo Fisher, Halethorpe, MD) utilizing the manufacturer’s protocol. Real-Time quantitative Reverse Transcription-PCR (qRT-PCR) was performed utilizing TaqMan® Gene Expression Master Mix or Fast Advanced Master Mix and TaqMan® Gene Expression Assays (Applied Biosystems™ by Thermo Fisher, Halethorpe, MD) with probes specific for GAPDH (expression control) and Mx1 utilizing the manufacturer’s protocols. qRT-PCR ran with the StepOnePlus™ machine and software (Applied Biosystems™ by Thermo Fisher, Halethorpe, MD).

### Statistics and Data

Tumor size, immunologic, RT-PCR, and flow cytometric analyses were statistically tested by Student’s t-test if two groups or by the one-way ANOVA with Tukey’s multiple comparison test if more than two groups. Mouse survival studies were statistically tested by the log-rank test. Scatter plots were analyzed by simple linear regression. Tumor time courses were analyzed by Area Under the Curve calculations with 95% confidence intervals (CI), where non-overlapping CI’s were considered significantly different or with two-way ANOVA with Tukey’s multiple comparisons test. Microsoft® Excel (Microsoft Corp, Redmond, Wa) was used for database management. Prism 7 (GraphPad Software, Inc. San Diego, CA) was utilized for statistical analyses and figure creation. A significance level of α≤0.05 was set for all experiments.

## RESULTS

### Addition of Trp-2 to vaccine

Previous work from our laboratory has shown that the fusion of MIP3α to gp100 enhanced the immunogenicity and efficacy of a DNA vaccine^4^. In the current experiment, a 100 amino acid region of the melanoma-associated antigen tyrosinase-related protein 2 (Trp2) was added to the construct to create a two-antigen vaccine referred to as MIP3α-GpTrp or MGpTrp. The region of Trp2 includes the known class-I immunodominant epitope Trp2_180-188_^16^. Figure 1A shows a diagram of the expressed constructs used in this study.

**Figure 1:**
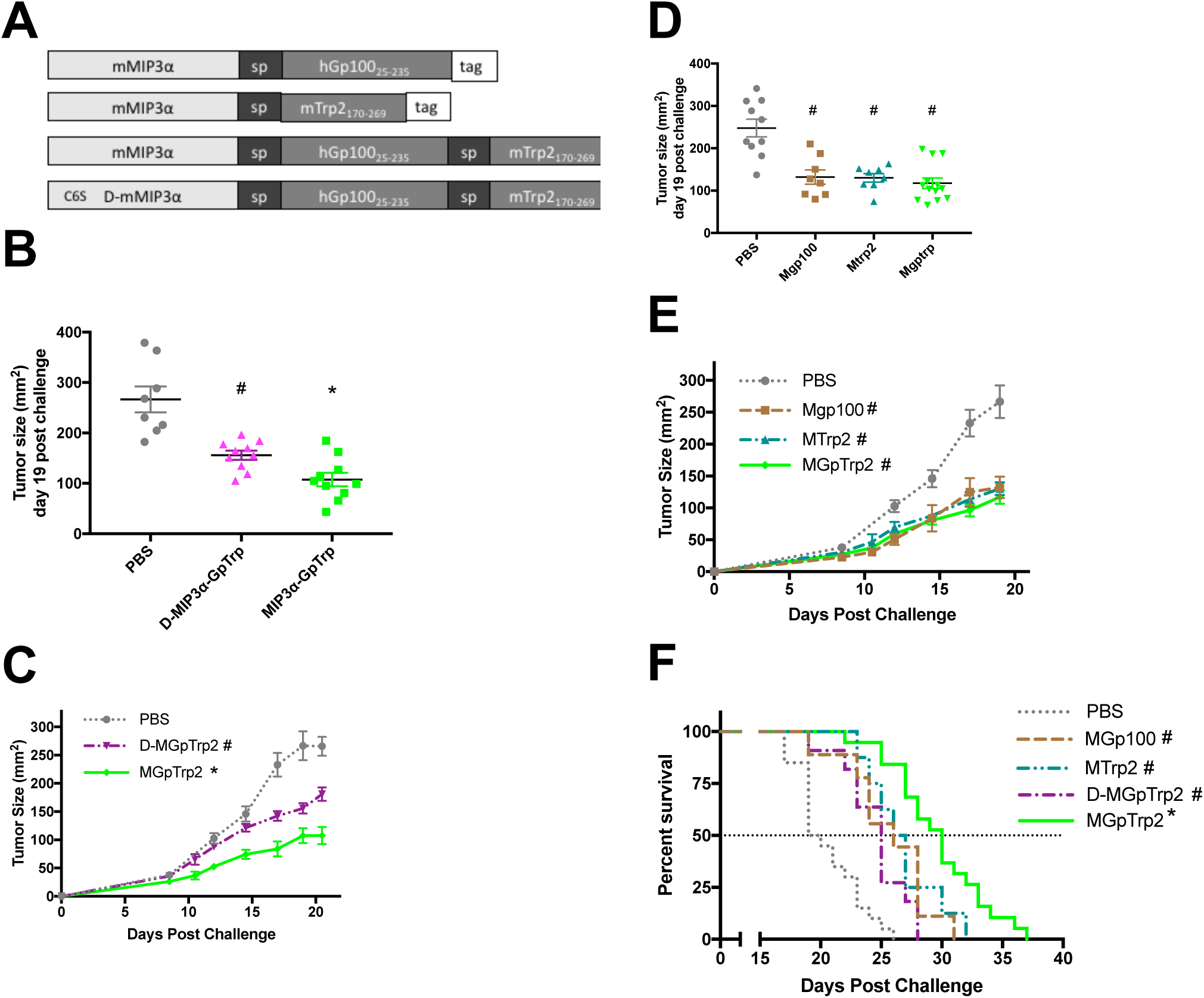
Vaccine Comparisons. A) Linear representations of expressed vaccine sequences within the vaccine plasmid. mMIP3α is full length and functional protein. D-mMIP3α contains a C6S mutation that renders MIP3α ineffective at targeting dendritic cells. “Sp” refers to a 14 amino acid spacer sequence. “Tag” refers to a 29 amino acid region including standard myc and histidine tags. Upstream of the construct is a secretion signal sequence from the mouse IP10 gene. For B-F, MGpTrp2 refers to vaccine with MIP3α fused to both Gp100 and Trp2 antigens. Mice were challenged at day 0 with 5×10^4^ B16F10 cells and vaccinated on days 5, 12, and 19 with 50μg plasmid B) Tumor size at day 19 post challenge and C) tumor growth time course from day 0 to day 21 comparing PBS mock vaccine, D-MIP3α-GpTrp2, and MIP3α-GpTrp2. D) Tumor size at day 19 post challenge and E) tumor growth time course from day 0 to day 19 comparing PBS mock vaccination, MGp100, MTrp2, and MGpTrp2. F) Kaplan-Meier survival analysis of all vaccine groups, assessed by log-rank test. Panels B-F show data combined from two to four independent experiments, n = 4-6 mice per group per experiment. Log(2) transformed tumor size data tested for significance by one-way anova with Tukey’s multiple comparison test. Tumor growth tested by Area Under the Curve calculations with non-overlapping 95% confidence intervals. Outliers more than two standard deviations from the mean were excluded from the dataset. #p<0.05 to negative control; *p<0.05 compared to all groups. Error bars denote estimate of standard error of the mean.

First, the comparative anti-melanoma efficacy of the MGpTrp double antigen vaccine compared to MGp100 and MTrp2 single antigen vaccines was tested. Three vaccinations were administered in one-week intervals beginning day five post B16F10 challenge, a point at which tumors were palpable at the inoculation site. At day 19 post challenge, all three vaccines had similar efficacy with percent reduction of tumor size between 47 and 53% compared to the PBS control (p<0.0001 vs PBS; Figure 1D). The early tumor growth patterns also showed that the three vaccines had similar efficacies up through day 19 post challenge (Figure 1E). However, prolonged efficacy in the MGpTrp2 group resulted in enhanced survival compared to MGp100 (p=0.01) and MTrp2 (p=0.04).

The next experiment addressed the efficacy of the MGpTrp vaccine compared to negative control and D-MIP3α-GpTrp control vaccine, in which the MIP3α component has been rendered inactive. Figure 1B shows that at day 19 post challenge, mice that received the MIP3α-GpTrp vaccine had a 63% reduction in average tumor size compared to the negative control (p<0.0001) and a 31% reduction compared to D-MIP3α-GpTrp vaccine (p<0.05). Tumor growth began to diverge between the three groups at day 10 post challenge (Figure 1C). Area Under the Curve (AUC) and survival analyses indicate that all three groups have significantly different tumor growth and mouse survival curves (Figure 1C and F).

### Enhancing the response with CpG adjuvant, IFNα, and Aza

Previous work from our laboratory provided evidence that neutralizing IL-10 at the tumor site could enhance therapeutic efficacy in a type-1 interferon pathway dependent manner^5^. It has also been shown that Aza can enhance the efficacy of IFNα therapy^10^, and that CpG oligonucleotides can enhance vaccine efficacy by mechanisms that include type-I interferon activity^10–12^. With this focus on type 1 interferon, a therapeutic protocol was designed with CpG adjuvanted MIP3α-GpTrp used in combination with Aza and a high-low dose series of IFNα treatments as outlined in Figure 2A. Henceforth, ‘vaccine’ will refer to the MGpTrp construct adjuvanted by CpG.

**Figure 2:**
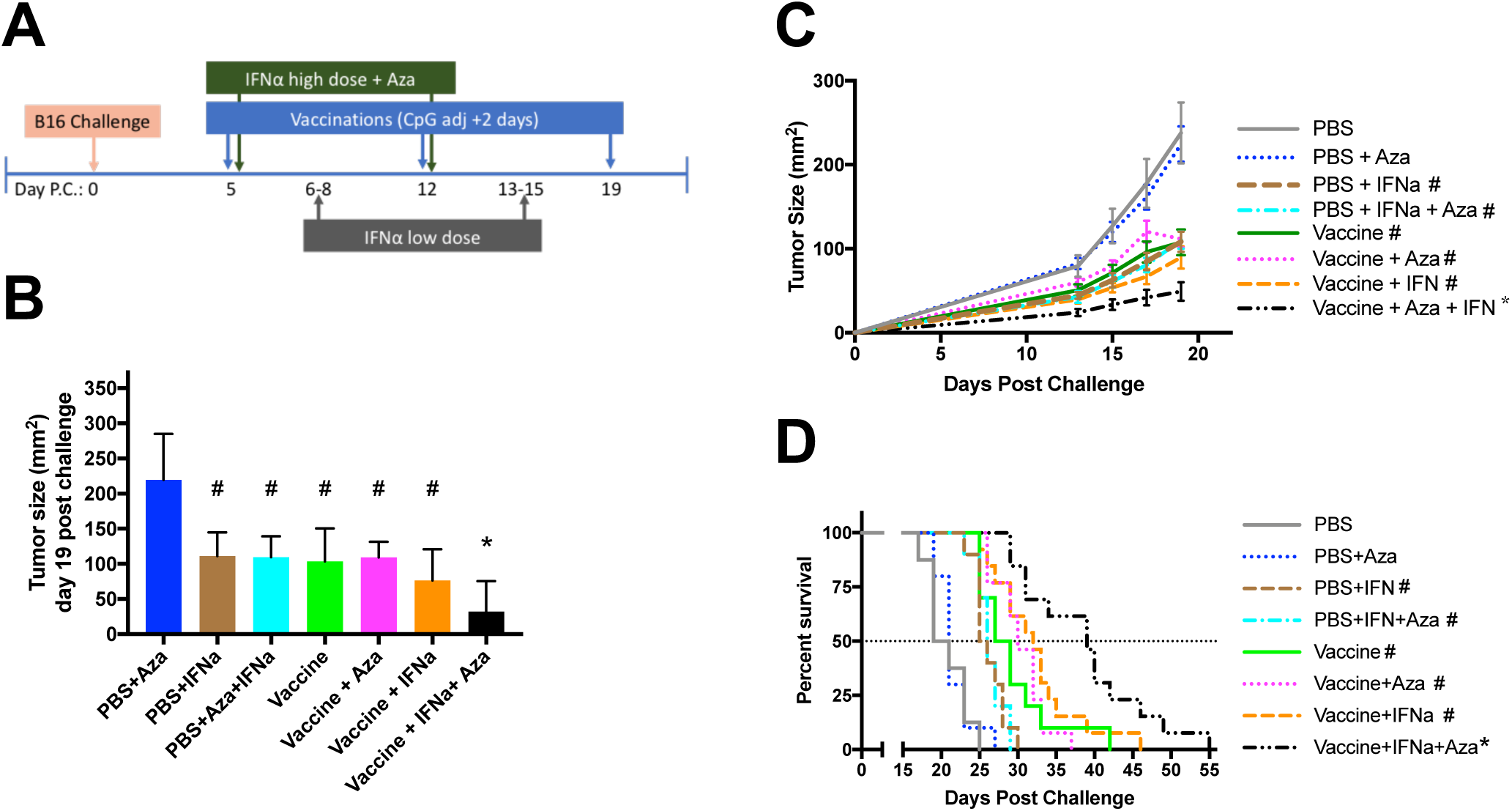
Addition of IFNα and Aza to Vaccine. Panel A shows the therapy schedule. At day 0 mice were challenged with 5×10^4^ B6F10 cells. Vaccine was composed of MIP3α-Gp100-Trp2 vaccine and given at 50μg/dose followed by i.m. electroporation. Two days after immunization, 50μg/dose CpG (ODN 2395) was given at immunization site. High dose IFNα (10,000 units) and low dose (1,000 units) were given intratumorally. Aza was given I.p. at 1mg/kg. B) Tumor sizes across groups at day 19 post challenge. PBS alone was excluded due to mice already removed from the group. Statistical analysis was performed on the log(2) transformed data values by one-way anova with Tukey’s multiple comparison test. C) Tumor size progression from day 0 to day 19. Significance was assessed by two-way ANOVA with Tukey’s Multiple Comparison Test. D) Kaplan-Meier Survival curve of all groups, tested by log-rank test. Panels B-D show the combined data from two independent experiments, n = 3-8 mice per group per experiment. Outliers more than two standard deviations from the mean were excluded from the dataset. On the graphs, statistical significance (α=0.05) is delineated by: # significant to PBS and/or PBS+Aza, * significant to all other groups. Error bars show estimated standard error of the mean.

On day 19 post tumor challenge, the tumor sizes of seven groups were analyzed, comparing the relative efficacies of all combinations of vaccine, Aza, and IFNα (Figure 2B). PBS only mice were excluded from analysis due to mice already reaching euthanasia thresholds. Judging by tumor growth in Figure 2C, the negative PBS vaccinated mice had similar tumor growth and size as the PBS vaccinated mice given Aza. Therefore, PBS+Aza mice are treated as the negative control comparator for figure 2B. Tumor growth over time up through day 21 post challenge was also monitored and tested (Figure 2C). In addition, mouse survival was assessed (Figure 2D). In all three of these figures, a similar pattern is seen. The use of IFNα provides an intermediate phenotype with or without Aza. The vaccine itself or with either IFNα or Aza but not both also provides an intermediate phenotype. However, when the vaccine is combined with both IFNα and Aza, superior anti-tumor efficacy is observed, with a 22-53% increase in median survival over the intermediate phenotype groups.

### Interferon stimulated gene expression

While IFNα and Aza are known to influence transcription of many genes^10–12,17^, of particular interest for these studies was a report indicating that Aza enhances upregulation of interferon stimulated genes such as Mx1^10^. To assess the transcription levels of Mx1 as a marker for such upregulation, two different therapeutic protocols were utilized. First, to assess transcription levels early in the therapy, a modified schedule was utilized as outlined in Figure 3A. For this schedule, the initiation of treatment was delayed until day 8 post challenge, but the intervals between treatments were left unchanged. This approach ensured tumors of adequate size for analysis at a timepoint 3 days after the first full round of treatment. Second, a late timepoint analysis utilized the standard schedule but with tumor harvested for analysis one week after the final vaccination (Figure 3A).

**Figure 3:**
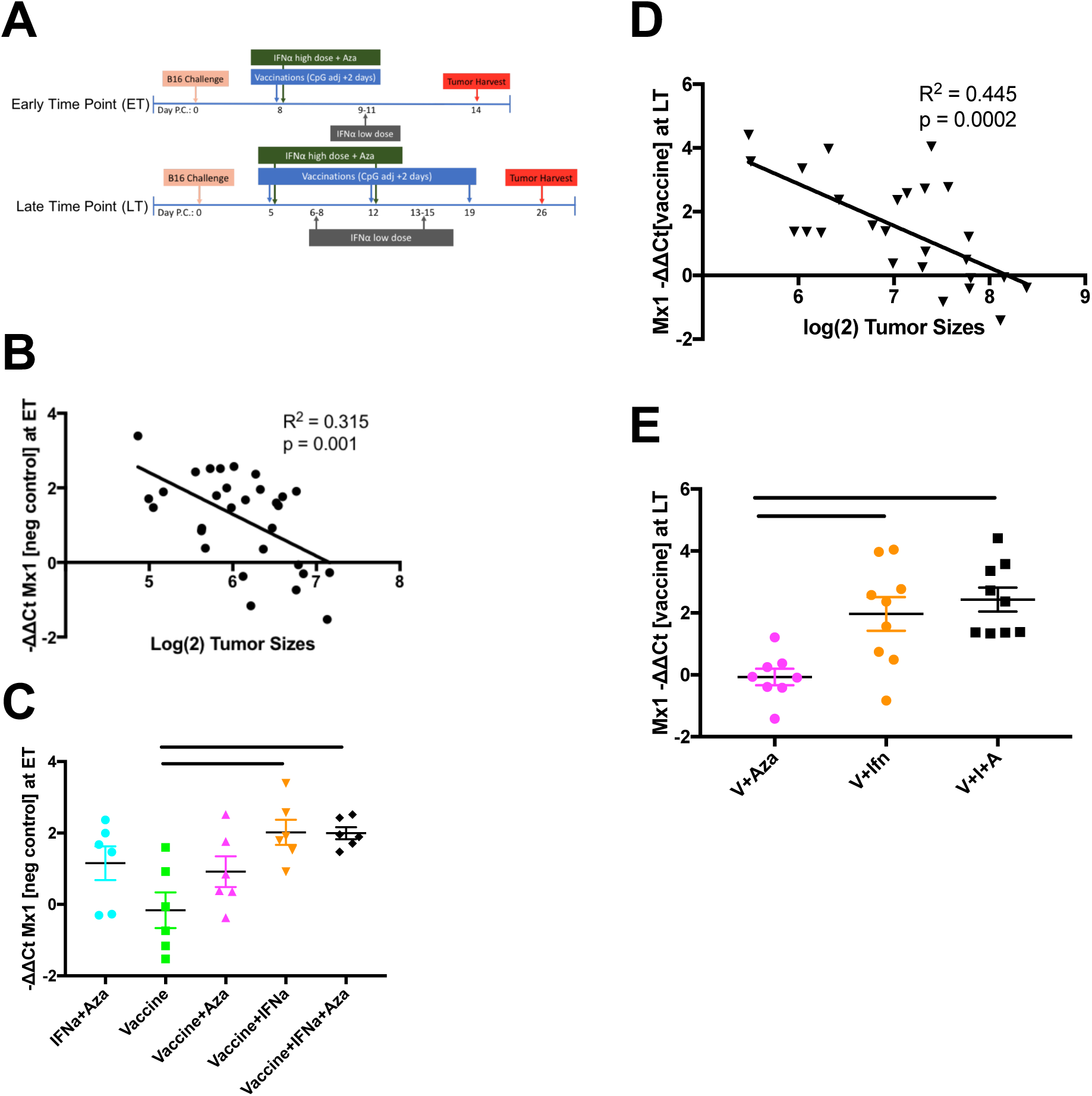
qRT-PCR analysis of gene expression. Panel A outlines the early and late time point therapy schedules. For analysis, ΔCt is calculated by subtracting the gene of interest Ct value from housekeeping gene GAPDH for each sample. ΔΔCt is calculated by subtracting ΔCt values from gene of interest to the ΔCt value of either negative control at the early timepoint or vaccine-only at the late time point. Panels B and D show the overall correlation between Mx1 -ΔΔCt values and tumor size at early (B) and late (D) time points. Panels C and E show the comparison across groups of Mx1 -ΔΔCt values at the early and late time points respectively. Panel data represent two independent experiments, n=3-5 mice per group per experiment. Scatterplots were tested by simple linear regression, with R^2^ and p values noted on the graphs. Grouped analyses were tested by Anova with Tukey’s multiple comparison test, with significant comparisons marked by bars. Error bars denote estimate of the standard error of the mean.

At the early timepoint, expression of Mx1 was negatively correlated with tumor size, with higher Mx1 transcription levels related to smaller tumors (Figure 3B). At this time point there was little difference in Mx1 expression between vaccine and the negative control (Figure 3C). IFN+Aza did not uniformly produce enhanced transcriptional levels of Mx1, resulting in a non-significant difference when compared to vaccine (p=0.21). Vaccine+Aza resulted in an intermediate level of Mx1 expression. Both vaccine+IFNα and vaccine+IFNα+Aza had significantly higher expression of Mx1 as compared to the vaccine alone. However, Aza did not further enhance Mx1 expression above levels measured in the vaccine+IFNα group. Notably, the early time point results remained consistent at the late timepoint, which is 11 days after the last dose of IFNα. The expression of Mx1 remained significantly negatively correlated to tumor size (Figure 3D). At this timepoint, vaccine alone served as the comparator and vaccine+Aza has roughly equivalent Mx1 expression to vaccine alone at this point. Both vaccine+IFNα and vaccine+IFNα+Aza maintained significantly higher expression of Mx1 as compared to vaccine+Aza (p<0.01) but the expression levels were not different compared to each other.

### Tumor infiltrating lymphocyte analysis

To analyze how the treatment regimens were influencing cellular composition of the tumor microenvironment, tumors were harvested at two timepoints (Figure 4A). Figure 4B outlines the gating strategy of the data in figures 4 and 5. Figures 4C-D show that at the early timepoint, the vaccine group, when compared to the PBS negative control mice or mice receiving IFNα+Aza without vaccine, had significantly higher percentages and normalized numbers of CD8+ TILs that produce activation cytokines after stimulation with vaccine peptides (p<0.01 for all). This finding is consistent with our previously published data^4^. While the percentage of TILs that were CD3-CD8 double positive were no different between the groups (Figure 4E), the vaccinated mice had higher normalized numbers of CD8+ T-cells infiltrating the tumor (Figure 4F; p<0.01). The groups combining vaccine with IFNα and/or Aza did not have tumors large enough at this time point to be analyzed by flow cytometry.

**Figure 4:**
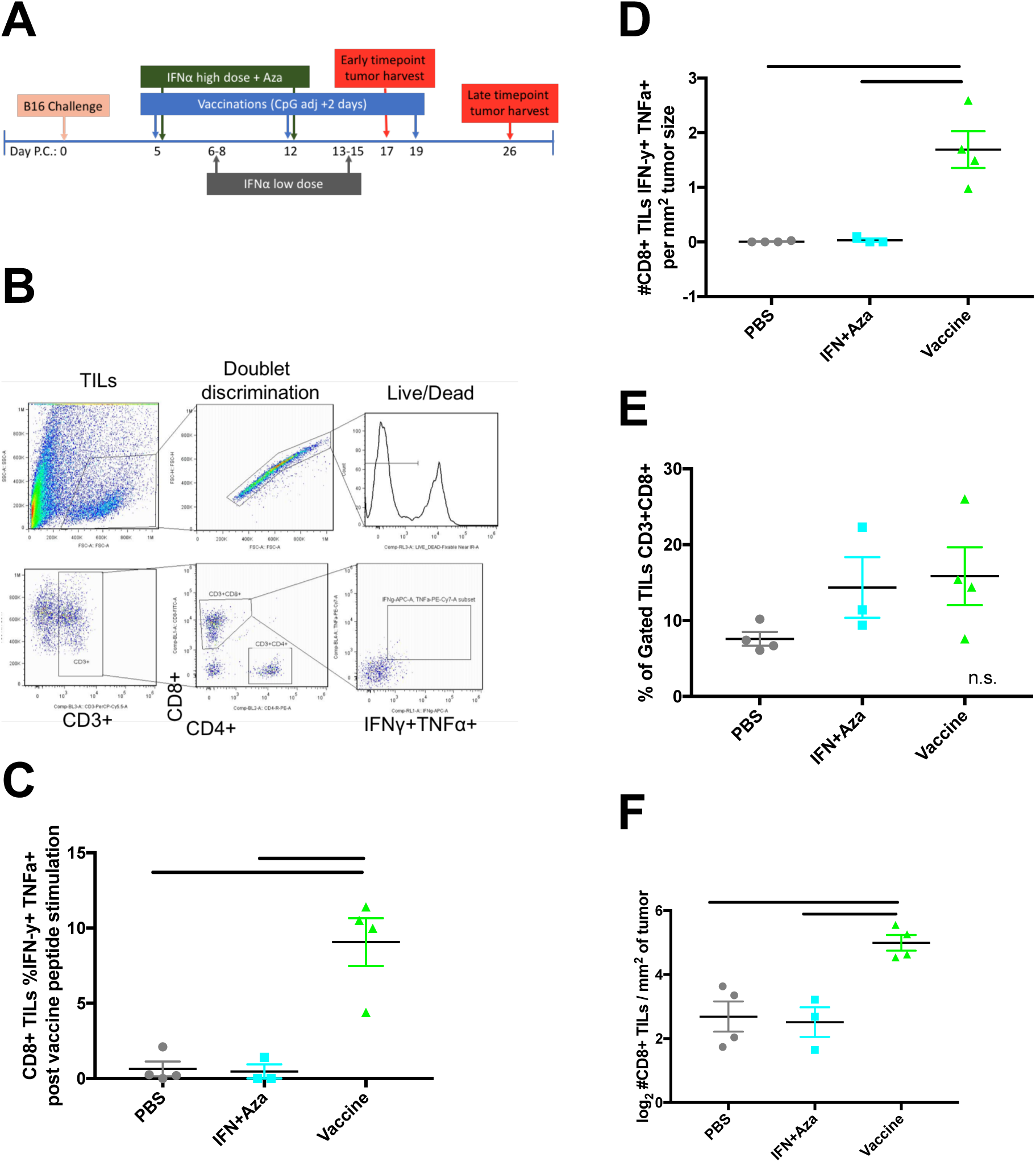
Early Time Point T-cell analysis. Panel A shows the therapy schedule. Vaccination and treatments are the same as outlined in Figure 2. This figure shows analysis of the early time point only. See Figure 5 for analysis of the late time point. Panel B shows the gating strategy. Leukocytes were gated by forward and side scatter from overall cells and were then screened by doublet and live/dead discrimination. TILs were determined by CD3 positivity followed by CD4 or CD8 positivity. CD3+CD8+ cells were then gated to determine IFNγ and TNFα double positivity indicative of productively stimulated effector cells. Panel C shows the percentage of and Panel D shows the tumor-size normalized numbers of CD3+CD8+ TILs that were IFNγ and TNFα double positive after stimulation with vaccine peptides. Panel E shows percentage of and Panel F shows the tumor-size normalized numbers of gated TILs that were CD3+CD8+. Data represents one experiment of n=3-4 mice per group. Significance tested by one-way Anova with Tukey’s multiple comparisons test, with significance noted by bars between groups. Error bars are representative of estimate of the standard error of the mean.

**Figure 5:**
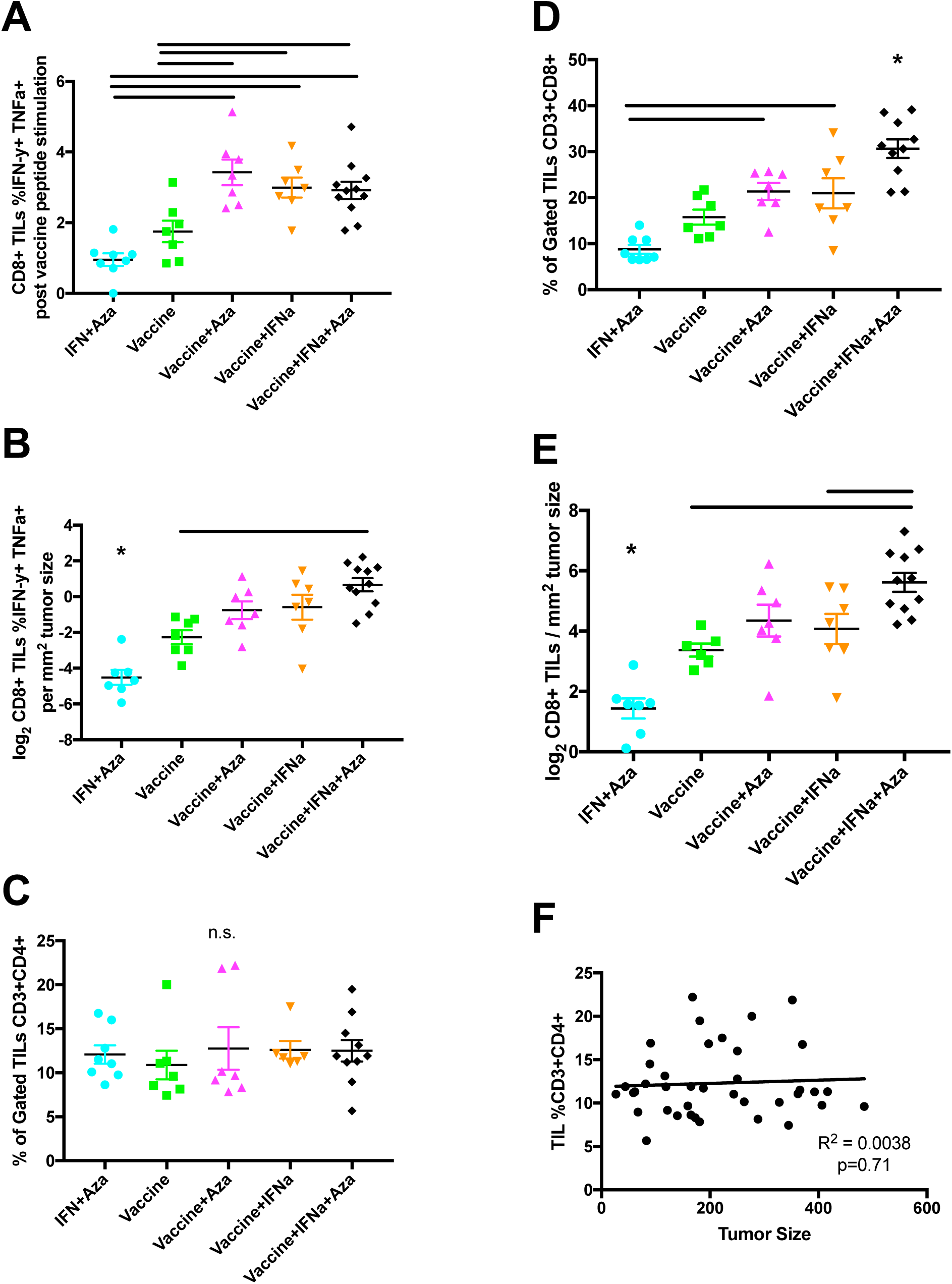
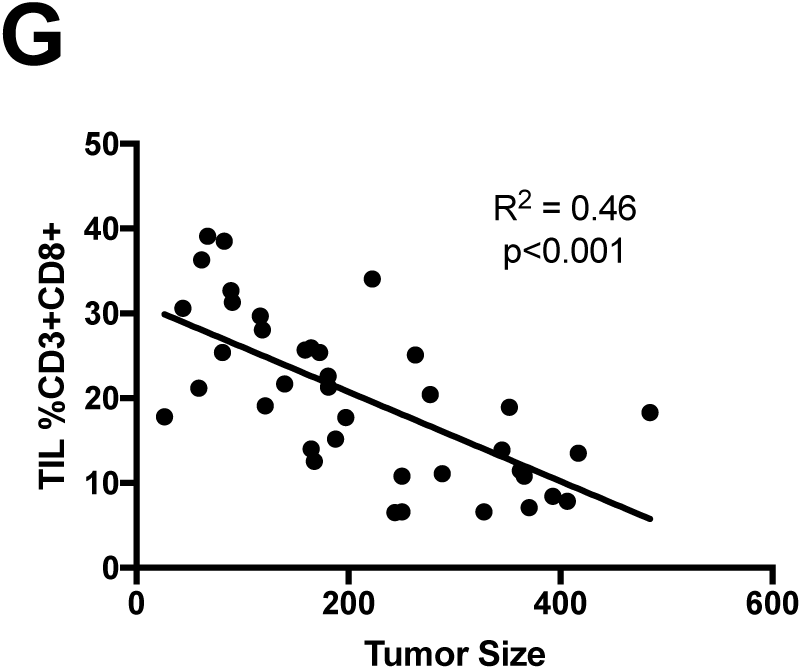
Late Time-Point T-cell analysis. Therapy schedule and gating strategy as outlined in figure 4. Data in this figure represents collection at the later time point. Panels A and B show the percentage and tumor-size normalized numbers respectively of CD3+CD8+ TILs that were successfully stimulated by vaccine peptides. Panel C shows the percentage of gated TILs that are CD3+CD4+. Panels D and E show the percentage and tumor-size normalized numbers respectively of gated TILs that are CD3+CD8+. Panels F and G show the scatterplots and correlations of mice from all groups comparing gated TILs that were CD3+CD4+ or CD3+CD8+ respectively to measured tumor size. All panels represent two to three independent experiments with n = 3-4 mice per group per experiment. Panels A-E were assessed by one-way Anova with Tukey’s multiple comparison test. Significance was annotated by bars between two groups or by an asterisk where the group is significantly different from all other groups. Panels F and G were tested by simple linear regression, with R^2^ and p values noted in the panel. Error bars denote estimate for the standard error of the mean. Outliers more than two standard deviations from the mean were removed from the dataset.

Panels of Figure 5 refer to the late time point tumor harvest. Figure 5A shows that after stimulation with vaccine peptide the percentage of CD8+ TILs that produce activation cytokines is elevated in the vaccine+Aza, vaccine+IFNα, and vaccine+IFNα+Aza groups as compared to IFNα+Aza (p<0.0001) and vaccine alone (p<0.05) groups. The three groups with elevated levels show no difference between them. However, when total vaccine-reactive CD8+ TILs were normalized for tumor size, the vaccine+IFNα+Aza group is the only one with significantly elevated levels as compared to vaccine alone (Figure 5B; p<0.01). Vaccine+Aza and Vaccine+IFNα have intermediate phenotypes. IFNα+Aza with no vaccination provides a negative baseline for the assay that is significantly different from all groups.

The overall TIL fraction was also analyzed. Figure 5C shows that by percentage there was no difference between groups with regards to CD3+CD4+ TILs. In contrast, the percentage of TILs that are CD3+CD8+ were markedly different (Figure 5D). Average percent levels of CD3+CD8+ TILs in vaccine+IFNα+Aza were elevated compared to all other groups. Vaccine+Aza and vaccine+IFNα groups had higher CD3+CD8+ TIL percentage than the IFNα+Aza group (p<0.01), but were not substantially higher than vaccine alone. Similar relationships are seen when estimating the total CD3+CD8+ TILs adjusted for tumor size (Figure 5E). Although not significantly higher as compared to vaccine+Aza by this measure, the vaccine+IFNα+Aza group has higher numbers of CD3+CD8+ TILs as compared to IFNα+Aza, vaccine alone, and vaccine+IFNα groups.

The correlations of tumor size and TIL percentage across groups were also analyzed. Figure 5F shows that the percentage of TILs that are CD3+CD4+ does not correlate with tumor size. In contrast, the percentage of TILs that are CD3+CD8+ does significantly negatively correlate with tumor size, with a higher CD3+CD8+ percentage correlating with lower tumor size (Figure 5G).

### Systemic administration of IFNα

One potential weakness of this treatment regimen is the administration of IFNα locally at the tumor site, which may not be feasible in the clinical setting. A study comparing intratumoral (i.t.) administration of IFNα to systemic intramuscular (i.m.) administration of IFNα was conducted using the standard schedule as outlined in Figure 2A. At day 19 post challenge, i.t. and i.m. administrations of IFNα produced equivalent tumor sizes when combined with vaccine and Aza (Figure 6A). Both had significantly reduced tumor size compared to the vaccine+Aza group (47-51% reduction, p<0.05). Similarly, i.t. and i.m. administrations of IFNα with vaccine and Aza provided nearly identical growth curves (Figure 6B). This pattern remained consistent with survival analysis (Figure 6C). Both i.t. and i.m. administrations of IFNα combined with vaccine and Aza had significantly enhanced survival as compared to the vaccine+Aza group (p<0.01).

**Figure 6:**
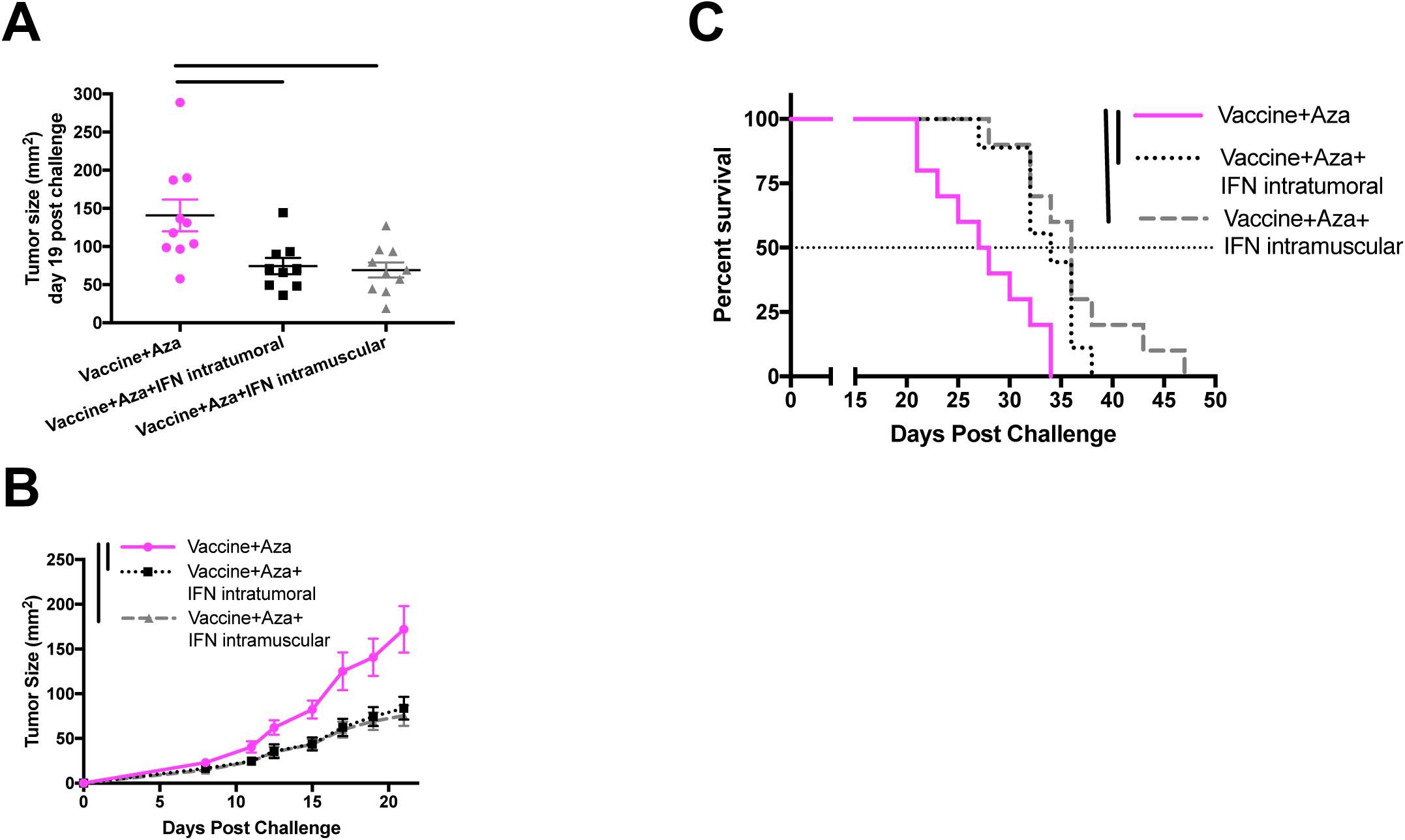
Intratumoral versus Intramuscular administration of IFNα. MIP3α-Gp100-Trp2 was used as the vaccine, with ODN2395 CpG adjuvant given two days later. Schedule of treatment is the same as Figure 2a. A) Tumor Sizes at day 19 post challenge. Data were tested by one-way Anova with Tukey’s multiple comparison test. B) Tumor growth time course through day 21 post challenge. Area Under the Curve statistics were calculated, and interactions were considered significant if 95% confidence intervals did not overlap. C) Kaplan-Meier survival curve as assessed by log-rank test. Panels A-C show combined data from two independent experiments, n = 3-7 mice per group per experiment. Statistical significance designated by bars between groups. Error bars show estimated standard error of the mean.

## DISCUSSION

In this study, we first provided evidence that the addition of a second antigen in the vaccine improved mouse survival. Vaccines with multiple antigens run the risk of antigenic competition^18^. However, many studies show multi-epitope vaccines with enhanced efficacy correlating with vaccine antigenic valency^14,15^, and in clinical trials, objective response rates of cancer vaccines were improved with multi-peptide strategies ^13^. The MIP3α-antigen vaccine platform had not yet been tested with multiple protein targets. Tyrosinase-related protein 2 (Trp2) is a melanoma-associated antigen similar to gp100 in that it has been successful in developing vaccine-derived anti-tumor immunity in preclinical models, and it has a known immunogenic MHC-I epitope (amino acids 180-188) ^19–22^. Additionally, use of a peptide of longer length has been shown to be enhance efficacy^23^, and therefore a 100 amino acid region of Trp2, including 180-188, was added to the MIP3α-gp100 construct to form a MIP3α-Gp100-Trp2 vaccine. The data showed that MIP3α continued to enhance the efficacy of a multi-epitope vaccine and that inclusion of both Gp100 and Trp2 antigenic regions together provided superior survival over either single antigen vaccine. The additive increase in efficacy shows that this platform is amenable to multi-epitope strategies. The addition of more tumor associated antigens and/or neoantigens as well as the relative immunogenicity of each antigen will be the subject of future studies.

We have shown previously that neutralizing IL-10 modestly enhanced vaccine therapeutic efficacy, an effect that was mediated through Type 1 interferon pathways^5^. IFNα is known to provide direct therapeutic efficacy via its ability to modulate anti-tumor immunity^24^, and it has also been studied as a potential vaccine adjuvant^25,26^. Furthermore, IFNα-2b therapy has shown clinical benefit^27^ and remains a part of the standard of care for certain melanoma patients^28,29^. To further enhance the potential effect of type 1 interferons in tumor control, we supplemented our vaccine regimen with an adjuvant known to elicit type 1 IFN activity. Synthetic CpG class C oligodeoxynucleotides act as danger signals that interact with toll-like receptor 9 on B-cells and plasmacytoid dendritic cells, thereby initiating immune signaling cascades that result in robust type-I interferon responses^30^. Studies have shown that the addition of such CpG based adjuvants can enhance the efficacy of DNA-based vaccines^31,32^.

A report by Lucarini *et al*. has described a novel combination therapy of type I interferons and 5-Aza-2’-Deoxycitidine (Dacogen®) that has improved efficacy against B16F10 mouse melanoma^10^. This report attributed the enhanced efficacy of this protocol to the ability of Aza to prevent inactivation of ISG’s attributable to gene silencing resulting from DNA methylation. Aza, has been utilized for decades for the therapy of leukemias^33^ and is currently in clinical trials for the treatment of solid tumors^34^. Aza acts as a DNA demethylating agent by inhibiting the DNA methyltransferase enzymes, ^35^ which play a role in epigenetic regulation of genetic transcription. Dysregulation of DNA methylation is a characteristic of cancers and promotes cancer cell survival and adaptation^36^ and is also known to affect melanoma progression and metastasis^37^. The data in figure 2 validates our primary hypothesis that Aza and IFNα would enhance anti-tumor efficacy of the MIP3α-GpTrp vaccine. The combination of the adjuvanted vaccine with recombinant IFNα and Aza provided superior anti-tumor efficacy over any other single or double agent therapy. Mouse median survival in particular almost doubled (95% increase) between negative control and triple agent therapy.

However, data from our studies did not support a finding that Aza was acting by enhancing or extending expression of ISG’s. We observe upregulated Mx1 expression when IFNα is added to the vaccine therapy, but that effect was not further enhanced by Aza. The difference from the Lucarini study might be due to a difference between our two experimental designs. They utilized five consecutive immunizations of 5×10^4^ U of IFNα^10^, whereas our study utilized a series of one dose at 1×10^4^ U followed by three doses of 1×10^3^ U, a more clinically realistic schedule.

Beyond their potential effect on ISG’s, the combination of Aza and type-I interferons in the B16F10 melanoma model has been reported to produce an increased percentage of CD8+ tumor infiltrating lymphocytes^10^. The data from our cellular infiltrate analysis were consistent with these findings. The overall CD8+ TIL infiltrate was enhanced. While the vaccine specific TILs were not selectively enhanced by percentage, the overall increase in TILs led to more vaccine-specific CD8+ TILs by number. A likely mechanism of the enhanced CD8+TIL infiltration could be due to increased intratumoral production of proinflammatory cytokines and chemokines and/or reduction of regulatory immune cells^10^. This mechanism will be further analyzed in future studies. Importantly, mice vaccinated without additional treatment developed a strong CD8+ T-cell response early that was lost later, whereas the groups including vaccine and IFNα still maintained a significant vaccine-specific CD8+ T-cell response at the later time point.

## CONCLUSION

The MIP3α-antigen vaccine platform has shown promise as a unique approach to cancer vaccination. Our results show that the platform is amenable to multi-antigen vaccine designs. Further, the vaccine efficacy can be greatly enhanced by the addition of agents affecting the type-I interferon pathway. The data indicate that the synergy seen in this study is due to the enhancement of inflammation induced by IFNα given either systemically or locally, leading to prolonged T-cell effector function, combined with the higher numbers of CD8+ TILs induced by the Aza and IFNα combination. This combination allows vaccine-specific CD8+ T-cells to provide anti-tumor efficacy more robustly and for a more sustained period of time. Further study is needed to fully understand the mechanisms underlying this enhanced efficacy, but the results achieved by this combination of agents in this preclinical model system defines an approach that merits testing in a clinical setting.

## LIST OF ABBREVIATIONS

- Aza: 5-Aza-2’-Deoxycytdine
- IFNα: Interferon alpha
- TIL: Tumor Infiltrating Lymphocyte
- MIP-3α: Macrophage Inflammatory Protein-3 alpha
- MGp100: Vaccine fusing MIP-3α to antigenic region of Gp100
- Trp2: Tyrosinase-related protein 2
- MtTrp2: Vaccine fusing MIP-3α to antigenic region of Trp2
- MGpTrp2: Vaccine fusing MIP-3α to antigenic regions of Gp100 and Trp2
- i.p.: intra-peritoneal administration
- i.m.: Intra-muscular administration
- i.t.: Intra-tumoral administration
- qRT-PCR: quantitative reverse-transcriptase polymerase chain reaction
- CCL20: C-C motif ligand 20
- CCR6: C-C Receptor 6
- iDC: Immature Dendritic Cells
- D-MIP3α: Defective version of MIP3α
- IL-10: Interleukin 10
- ISG: Interferon Stimulated Gene
- AUC: Area under the curve

